# Electrode-Tethered DNA: Interfacial Mechanics and Switching Dynamics from Ensemble-Level Atomistic Simulations

**DOI:** 10.64898/2026.09.29.755399

**Authors:** Yechan Noh, Huibin Chang, Kaiyu X. Fu

## Abstract

Surface-tethered DNA, anchored to gold by the late 1990s, has matured into a versatile platform for biosensing, computing and actuation. Whereas the molecular behavior of DNA in bulk solution is well characterized, that of DNA at electrode interfaces remains far less understood. Here we investigate the mechanics and dynamics of electrode-tethered double-stranded DNA across surface chemistries, salt concentrations and applied potentials. Using GPU-accelerated all-atom molecular dynamics, we sample trajectory ensembles totaling 144 *µ*s over 620 simulations. The simulations reproduce key experimental observations yet uncover molecular behavior that had gone unnoticed despite nearly three decades of research. Ensemble statistics reveal a preferred orientation about the helical axis upon surface contact. Energetic analysis identifies a substantial role for Onsager–Samaras-type repulsion in opposing that contact. By reproducing experimental trends while resolving their molecular origins, these ensembles substantially narrow the longstanding simulation–experiment gap and provide a basis for engineering tethered-DNA devices.

## Introduction

DNA has been studied electrochemically for more than six decades.^1^ In 1997–1998, thiol attachment and alkanethiol passivation enabled controlled organization of DNA on gold.^2–4^ In 1998, applied potentials were shown to reversibly change the orientation of tethered DNA helices. ^5^ Since the late 2010s, surface-tethered DNA has found increasing use in molecular devices. More broadly, DNA-based devices now span sensing,^6–9^ nanomachines,^10–14^ computing^15,16^ and repeated information retrieval.^17,18^ At electrodes, controlled DNA motion can both report target binding^6^ and assist sensor resetting through the release of bound analytes.^19^ These functions depend on DNA configuration, motion and molecular contacts at the interface.

Tethering DNA to a surface constrains its geometry and exposes it to interactions with the substrate and surrounding electrolyte that often govern its behavior.^20,21^ Experiments have mapped the resulting responses in detail, from potential-driven switching between lying and standing states^22,23^ to the dependence of orientation on electrolyte conditions.^21^ These optical or electrochemical measurements reveal how tethered DNA behaves, but the underlying mechanisms are typically inferred by interpreting the signals through physical models.^20,24^ Despite decades of successful application, many of these mechanisms therefore remain to be verified at the molecular level.

Simulating such a system at atomic resolution has long been prohibitively expensive because an explicit model of the DNA, electrode, passivation layer, water and ions readily exceeds 100,000 atoms. Prior work has accordingly relied on simplified descriptions: Brownian dynamics^23^ and coarse-grained models^25^ reach the long timescales of tethered-DNA reorientation with reduced molecular detail. All-atom molecular dynamics (MD) has also been applied, but not at the ensemble scale needed to sample DNA motion broadly across conditions: it has instead resolved interfacial interactions within selected DNA–surface systems.^26–28^ Reorientation in tethered DNA, however, is stochastic and slow, requiring an ensemble of long trajectories for each combination of applied potential and electrolyte conditions.

Here we investigate the molecular basis of electrode-tethered double-stranded DNA (dsDNA) mechanics and motion through ensembles of GPU-accelerated all-atom MD simulations.^29^ We systematically compare surface chemistries, salt concentrations and electrode charges to determine how tethering modifies DNA elasticity and how interfacial interactions govern equilibrium orientation, voltage-driven lie-down and recovery after bias removal. The resulting ensembles test existing mechanistic interpretations and reveal previously unrecognized behavior. By reproducing established experimental trends while resolving their molecular origins, they narrow the gap between molecular simulation and experiment.

## Results and Discussion

### Structure and mechanics of electrode-tethered DNA

We first examine how electrode bias and salt concentration affect the structure and mechanical response of tethered dsDNA. The model system is 24 bp dsDNA attached through a C6 thiol linker to gold passivated with 6-mercapto-1-hexanol (MCH) in aqueous NaCl at 310 K (Fig. 1A). Concentrations refer to added NaCl: additional ions for charge compensation and double-layer formation vary with surface chemistry and bias (Methods, “System construction”: Supplementary Table S2). We characterize the structural response through end-to-end length and mean helical twist.

**Fig. 1.**
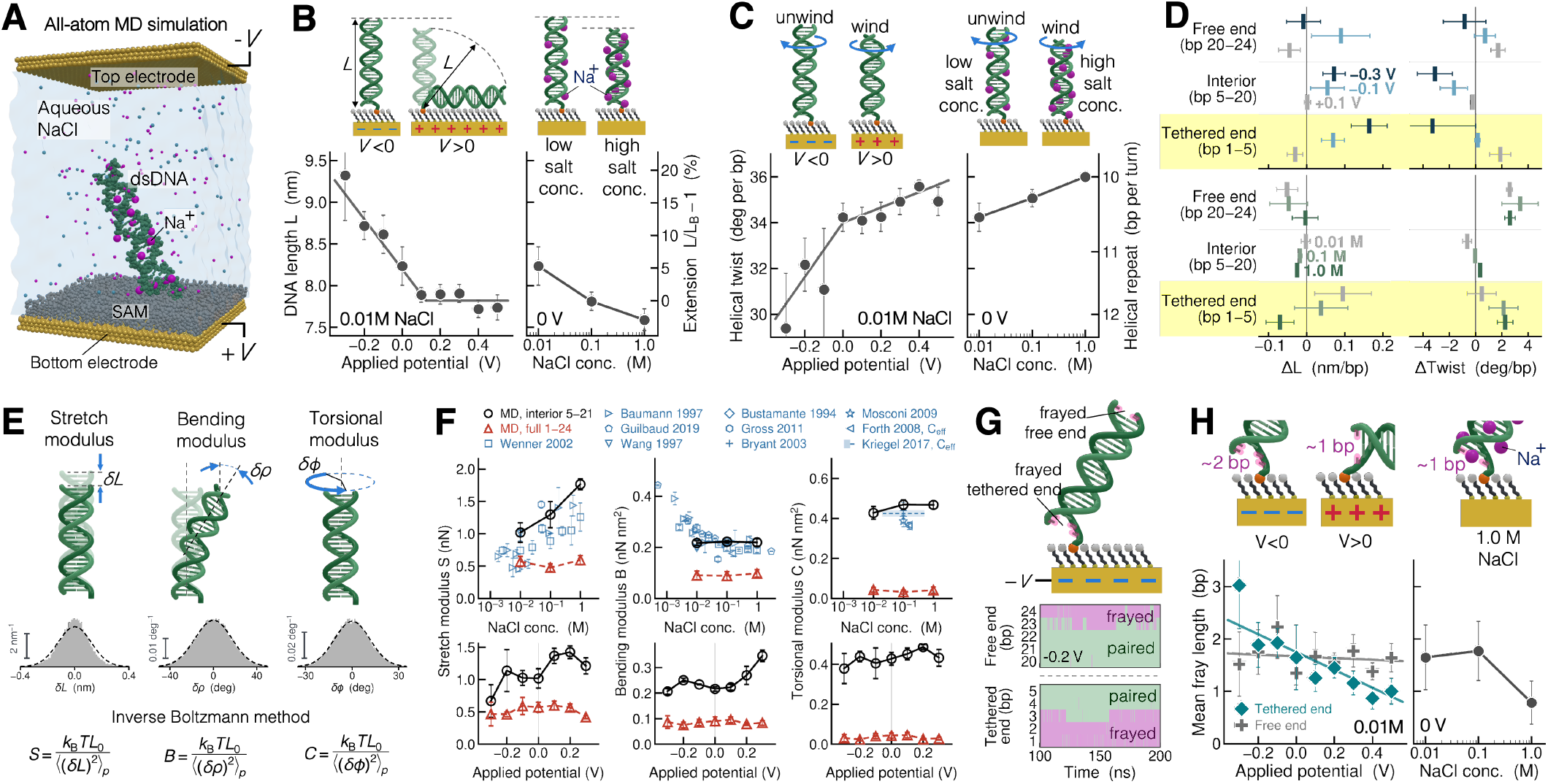
Structure and apparent mechanical properties of electrode-tethered ds-DNA. **A**, Simulation system with 24 bp dsDNA tethered through a C6 thiol linker to MCH-passivated gold. **B**,**C**, End-to-end length (B) and helical twist (C): *L*_*B*_ denotes the ideal B-form end-to-end length. **D**, Regional changes in base-pair spacing and twist relative to the reference averaged over +0.2 to +0.5 V at 0.01 M NaCl: potential series (top), salt series (bottom). **E**, Fluctuation distributions at zero bias and 0.1 M NaCl: dashed curves are Gaussian fits. In the expressions, *L*_0_ is the fragment reference length and subscript *p* denotes conditional variance. **F**, Apparent moduli versus NaCl concentration (top) and applied potential (bottom), with published measurements.^32–37,46–49^ The blue band denotes the experimental high-force effective torsional modulus. **G**, Base-pairing states in one trajectory at 0.01 M NaCl. **H**, Mean frayed length at the tethered and free ends (left) and at the tethered end (right). In B, C, D, F and H, potential series vary bias at fixed 0.01 M NaCl: salt series vary NaCl concentration at fixed zero bias. All simulations are at 310 K. Simulation points and error bars denote replicate means and SEM, respectively: experimental uncertainty conventions are given in Supplementary Table S6. Curves in B, C, F and H are guides to the eye.

At zero bias, raising NaCl concentration from 0.01 to 1.0 M shortens the dsDNA by approximately 7.8% and increases its mean helical twist angle by 5.2% (Fig. 1B,C). The increase in twist is qualitatively consistent with measurements on DNA in solution^30,31^ (Supplementary Table S5). Negative electrode bias produces the opposite structural response: at 0.01 M NaCl, the mean end-to-end length is approximately 18% greater at −0.3 V than at +0.1 V, while the mean helical twist decreases.

These changes are not distributed uniformly along the helix. The largest voltage-induced changes in base-pair spacing and local twist occur near the electrode attachment (Fig. 1D). At −0.3 V, the spacing increase relative to the positive-bias reference is more than twice as large near the tether as in the interior. This uneven structural response raises the question of whether elastic constants measured for DNA in solution can represent the tethered molecule.

To assess the mechanical consequences of these structural changes, we extract stretching, bending and torsional stiffness from conformational fluctuations using the inverse Boltzmann method (Fig. 1E). These moduli are apparent because the fluctuations include structural disruption as well as elastic deformation. At 0.1 M and zero bias, fragments sampled across bp 1–24, including both ends, yield a stretch modulus *S* of approximately 0.475 nN, a bending modulus *B* of 0.0884 nN nm^2^ and a torsional modulus *C* of 0.0317 nN nm^2^ (Fig. 1F). Single-molecule measurements on DNA in solution at approximately 0.05–0.186 M monovalent salt span approximately 0.88–1.45 nN, 0.15– 0.22 nN nm^2^ and 0.39–0.44 nN nm^2^ for *S, B* and *C*, respectively.^32–37^ The apparent moduli are therefore approximately 46–67%, 41–60% and 92–93% lower, respectively, with the greatest reduction in torsional stiffness. However, restricting fragment sampling to the interior, bp 5– 21, with the same frames and fragment lengths gives *S* ≈ 1.294 nN, *B* ≈ 0.222 nN nm^2^ and *C* ≈ 0.469 nN nm^2^, close to experimental values (Supplementary Table S5). The dsDNA interior thus has elastic moduli close to solution measurements, whereas including the terminal regions substantially lowers the apparent stiffness.

End fraying provides a physical explanation for this lower apparent stiffness: loss of base pairing reduces local dsDNA rigidity. Near the tether, this loss becomes more pronounced under negative bias, with repeated opening at −0.2 V (Fig. 1G). The mean frayed length at this end increases from approximately one base pair under positive bias to three at −0.3 V, whereas the free end responds less strongly (Fig. 1H). Increasing salt suppresses tethered-end fraying, linking terminal stability to the interfacial ionic environment. The enhanced fraying under negative bias is qualitatively in line with experimental reports of voltage-induced denaturation of gold-immobilized DNA.^38^ DNA tethering thus not only imposes a geometrical constraint but also alters the apparent mechanical properties of dsDNA.

### Equilibrium orientation and its physical basis

In electrode-tethered DNA devices, the orientation of dsDNA governs its proximity to the electrode and its accessibility to molecules in solution. We compare equilibrium configurations at atomic resolution on net-neutral MCH with polar hydroxyl headgroups, anionic MHA, cationic AHT and bare gold at zero bias and 310 K in 0.01–1.0 M NaCl (Fig. 2A–D). Tilt *θ* is measured from the surface normal, 0° upright and 90° lying flat. On net-neutral MCH the dsDNA stands predominantly upright, with mean tilts of approximately 23°, 32° and 28° at 0.01, 0.1 and 1.0 M NaCl (Fig. 2C,D). The mean tilt also peaks at 0.1 M in an independent ensemble carrying a methylene blue (MB) label (Fig. 2D).

**Fig. 2.**
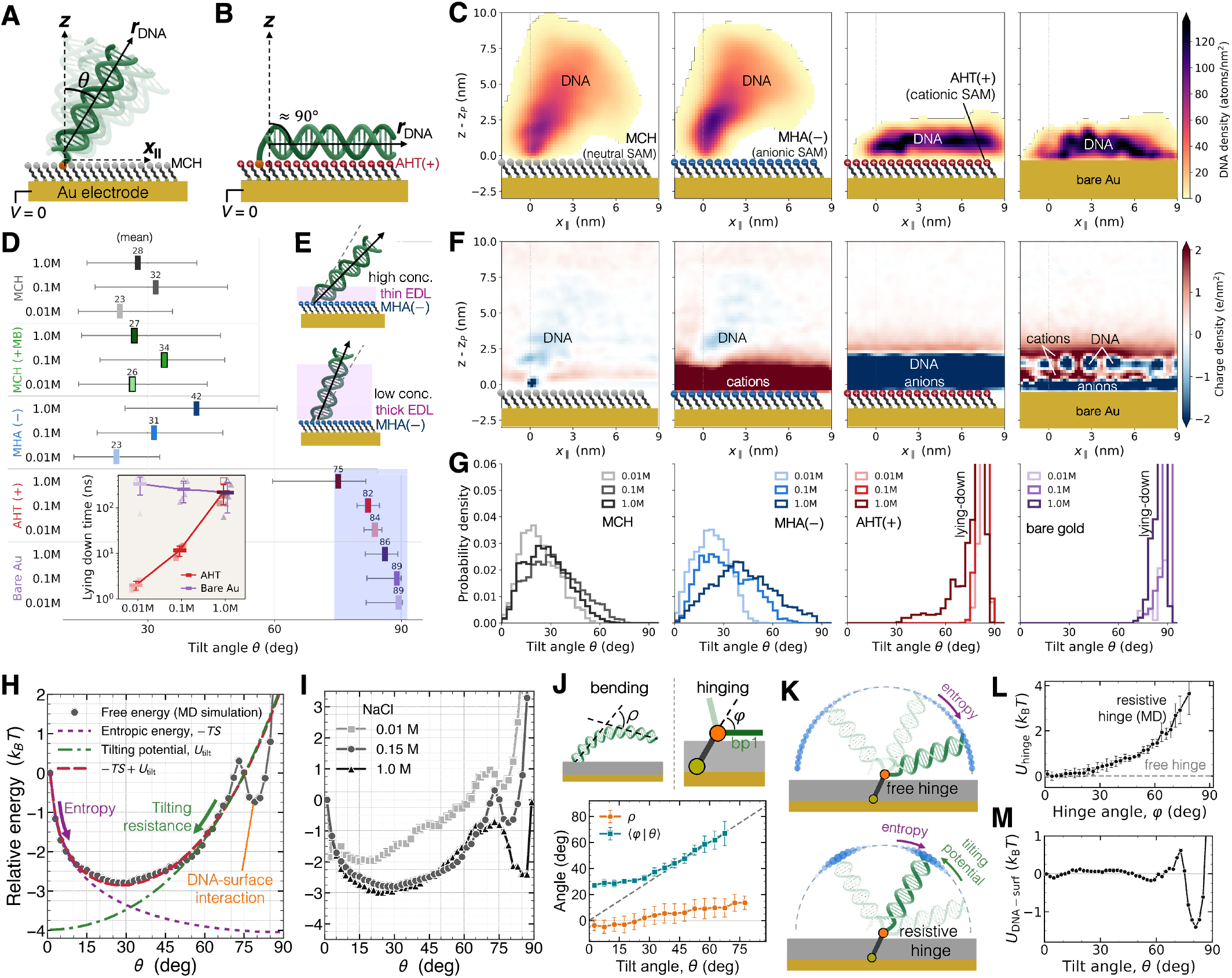
Surface-dependent orientation and resistance to reorientation of tethered dsDNA. **A**,**B**, Schematics defining tilt *θ* relative to the surface normal. **C**, DNA density maps in the molecular frame. **D**, Mean tilt across surface chemistries and salt concentrations: gray bars denote root-mean-square deviations above and below the mean. MB, methylene blue. Inset, mean±SD times to sustained *θ* ≥ 70°: open symbols mark observation limits for trajectories without lie-down. **E**, Schematic of salt-dependent ionic screening on MHA. **F**, Charge-density maps of ions and DNA phosphates, excluding monolayer charges: surface order as in C. **G**, Tilt distributions: AHT and bare-gold data in D and G describe adsorbed configurations. **H**, Orientational free energy and its decomposition on MCH. **I**, Salt dependence of the orientational free energy on MCH: quantitative interpretation of the 1.0 M curve is restricted to *θ* ≤ 70°. **J**, Bending angle *ρ* and joint angle *φ* versus overall tilt *θ* on MCH. **K**, Free- and resistive-hinge models. **L**, Effective joint potential on MCH. **M**, DNA–surface free-energy contribution on MCH. All panels describe zero electrode bias at 310 K: MHA and AHT have 10% ionized headgroups. C, F, J and L correspond to 0.1 M NaCl: H and M to 0.15 M. In C and F, *x*‖ follows the tilt azimuth and *z*_*P*_ is the anchor-phosphate height. Error bars in J and L denote estimated standard errors (Methods, “Free-energy landscape and joint geometry”).

The 10% ionized MHA and AHT monolayers carry −7.4 and +7.4 *µ*C cm^−2^, respectively, densities equivalent to nominal electrode biases of ∓0.37 V under the charge–voltage conversion of the model (Methods, “Applied potential”). On anionic MHA the mean tilt rises from approximately 23° to 42° between 0.01 and 1.0 M NaCl, as expected from screening of the DNA–surface repulsion (Fig. 2D,E). At 1.0 M, however, the dsDNA lies further over on anionic MHA than on net-neutral MCH, the opposite of what repulsion between the monolayer and the DNA backbone would suggest. Atomic-resolution analysis of the local ionic environment reveals a layer of accumulated cations above the MHA headgroups at 0.1 M (Fig. 2F). Attraction to these accumulated cations provides a physical explanation for the larger tilt at high salt.

On AHT and bare gold, in contrast, the dsDNA lies down spontaneously, reaching mean tilts of 75–84° and 86–89°, respectively (Fig. 2C,D,G), though not by the same route. On AHT the driving interaction is electrostatic and its range decreases with salt: raising NaCl from 0.01 to 1.0 M slows lie-down roughly a hundredfold (Fig. 2D, inset). At bare gold, van der Waals attraction acts only near the surface. The lie-down rate is therefore set by rotational diffusion that brings the dsDNA into this region, so no pronounced salt dependence is expected.

Tethered dsDNA is often idealized as a molecular pendulum, with tilting modeled as rotation about a free hinge.^6,20^ Free-hinge rotation is governed by angular orientational entropy, with a free-energy contribution given by

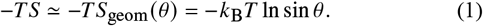

This entropic contribution drives DNA toward the surface because more orientational space is available at high tilt angles. However, our simulated mean tilt angles on MCH-passivated gold are approximately 23– 32°, consistent with the estimated tilt angle of 30° from neutron reflectivity measurements at 1 M NaCl.^3^ This upright orientation has previously been attributed to asymmetric attachment and steric interactions with the monolayer.^21^ However, the salt-dependent tilting resistance in our free energies is not explained by these mechanisms alone (Fig. 2H,I).

To examine the origin of this resistance, we distinguish joint rotation from duplex bending at atomic resolution. We find that DNA tilting involves mostly hinging and weak bending: the joint angle *φ* follows the overall tilt, whereas bending of the adjoining helix remains below approximately 15° even at *θ* = 75° (Fig. 2J). The effective joint potential increases by approximately 3 k^B^T across the sampled range, indicating a resistive rather than a free hinge (Fig. 2K,L). We attribute tilting resistance to bending energy at the joint, modest dsDNA bending, and ion and water rearrangement: their individual contributions remain unresolved. Based on the increasing stiffness with tilt (Fig. 2L), we approximate the combined resistance with a hyperbolic-cosine potential, which reduces to a harmonic hinge near the upright state and steepens at larger tilt,

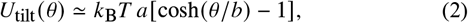

where *a* is a dimensionless amplitude and *b* an angular scale, with angles in radians (Methods, “Free-energy landscape and joint geometry”).

A further contribution arises from DNA–surface interactions (Fig. 2H,M). Equilibrium tilting at zero bias therefore reflects competition among orientational entropy, tilting resistance and DNA–surface interactions:

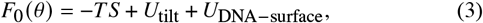

where the effective interaction terms include energetic and entropic contributions beyond angular orientational entropy.

On MCH, *U*_DNA−surface_ mainly acts at high tilt angles above approximately 70°. Within this effective interfacial contribution, favorable contacts involving the terminal hydroxyl groups compete with Onsager–Samaras-type repulsion.^39^ This repulsion arises because the surface restricts the counterion atmosphere surrounding negatively charged DNA. The stronger repulsive contribution at lower salt weakens contact stabilization, consistent with the shallower near-contact free-energy minimum at 0.01 than at 0.15 M NaCl (Fig. 2I). Our results reveal a previously unrecognized role for this repulsion in DNA– surface contact, even on a monolayer with no net charge at zero bias. Its consequences extend to lie-down and recovery dynamics, where it plays an important role in the formation and release of surface contact, as discussed below. The balance of attractive and repulsive DNA– surface interactions therefore influences dsDNA orientation even on net-neutral surfaces and can dominate it on charged passivation layers or under an applied electrode potential.

### Electrical actuation, ion dynamics and DNA lie-down orientation

We next examine DNA tilting driven by an applied electrode potential, the resulting switching dynamics, and the molecular contact DNA establishes with the electrode. On net-neutral MCH, positive electrode bias drives upright dsDNA toward the surface (Fig. 3A,B), consistent with earlier experiments.^21,22^ At 0.01 M NaCl, increasing the bias from +0.2 to +0.5 V shortens the Kaplan–Meier median lie-down time from 22.1 to 3.8 ns, nearly sixfold (Fig. 3L). Note that these lie-down times reflect DNA and ion response to an already-imposed electrode charge, whereas experimental switching times additionally include external-circuit charging and thus reflect coupled charging and DNA reorientation dynamics rather than DNA motion alone.^23^ Lie-down is also slower with TIP4P/2005 water than with TIP3P at the same nominal bias and salt concentration (Supplementary Fig. S1), further illustrating the model dependence of the response time.

**Fig. 3.**
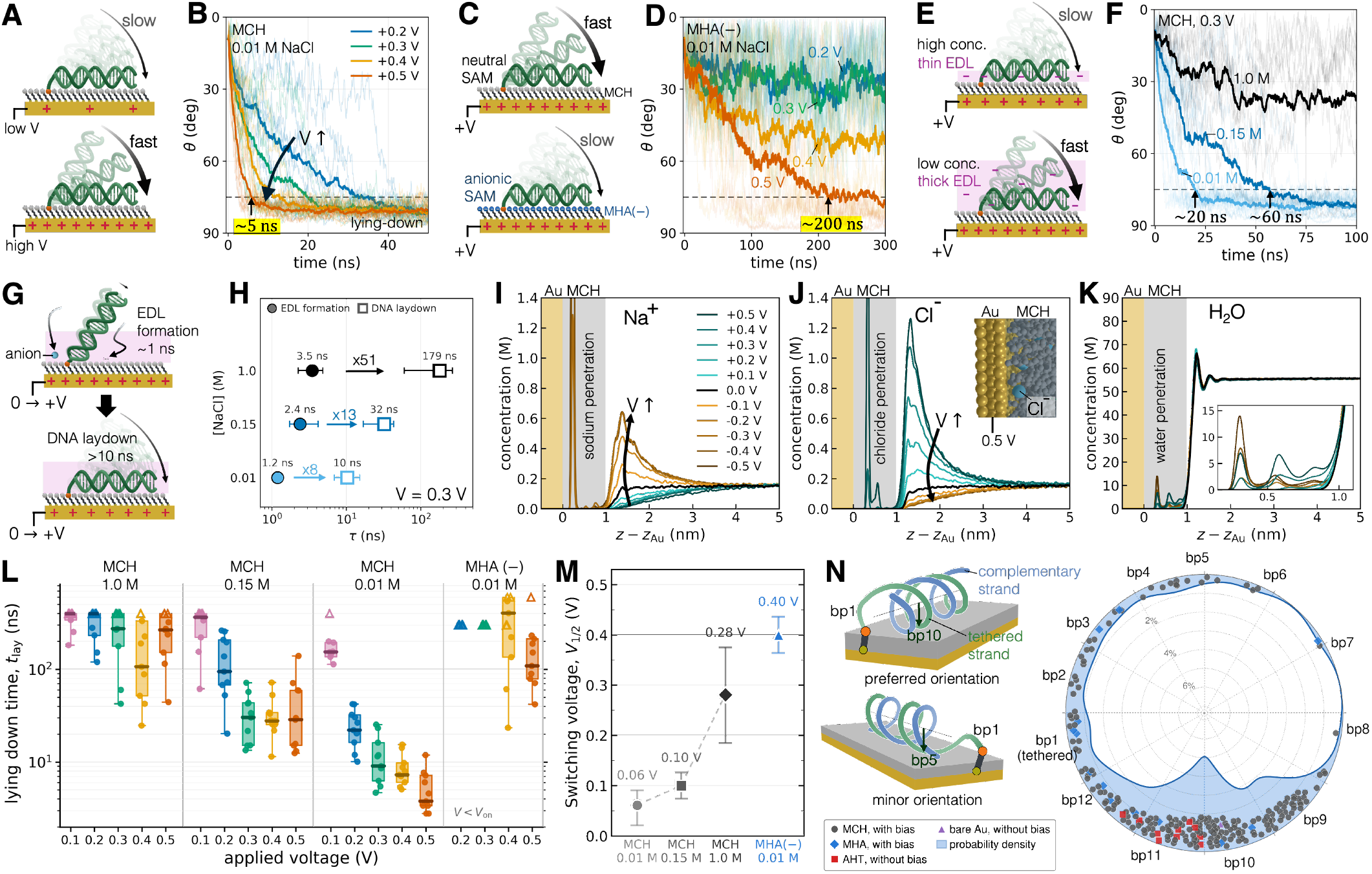
Electrical switching and surface contact of tethered dsDNA. **A**,**B**, Lie-down with varying bias on MCH at fixed 0.01 M NaCl. **C**,**D**, Corresponding voltage series on anionic MHA. **E**,**F**, Lie-down with varying NaCl concentration on MCH at fixed +0.3 V. Bold and faint traces in B, D and F denote ensemble means and individual trajectories: dashed lines mark the 75° lie-down threshold. **G**, Schematic of electric-double-layer (EDL) formation followed by DNA lie-down. **H**, EDL relaxation time constants (circles) and median observed lie-down times (squares) on MCH: error bars denote 68% intervals and interquartile ranges, respectively. **I– K**, Voltage-dependent Na^+^, Cl^−^ and water concentration profiles, respectively, for MCH-passivated gold without DNA at 0.15 M NaCl. **L**, Lie-down times across voltage, salt concentration and surface chemistry. Boxes show Kaplan–Meier medians and interquartile ranges: unreached values are shown at the observation limit. Circles mark observed events and whiskers span the observed event times. Open and filled triangles mark trajectories and conditions without observed lie-down, respectively. **M**, Fitted voltage for 50% lie-down within the observation windows, *V*_1/2_: error bars denote 95% conditional likelihood intervals. **N**, Orientation about the DNA helical axis upon surface contact, viewed from the tethered toward the free end. Each marker represents one trajectory (*n* = 223): base-pair labels indicate helical phases. Shading denotes probability per 5° window. All simulations are at 310 K.

Surface charge on the passivation layer substantially changes DNA tilting dynamics. On anionic MHA at 0.01 M, lie-down remains slow even under strong positive bias: at +0.5 V, the Kaplan–Meier median lie-down time is 109 ns, compared with 3.8 ns on MCH (Fig. 3C,D,L). The opposing surface charge can therefore delay tilting by imposing an energy barrier to lying down. Our simulations thus demonstrate the crucial role of surface charge in DNA tilting dynamics. Because surface charge depends on pH the interfacial *pK*_*a*_,^40,41^ these quantities can serve as important design parameters for tuning this response. The simulated voltages describe this model and should be interpreted and as trends rather than transferred directly to experiments.

Salt concentration also strongly changes the response time. At +0.3 V on MCH, the observed-event median increases from 10.4 to 32.2 and 179 ns at 0.01, 0.15 and 1.0 M NaCl, respectively, an approximately 17-fold slowing across this range (Fig. 3E,F,H). Lie-down occurs in all ten trajectories at each of the two lower salt concentrations, but in only five of ten at 1.0 M within the observation window. This trend follows the shortening of the electrostatic screening length at higher salt, which weakens the electrical attraction driving lie-down. In contrast, electric-double-layer (EDL) formation, monitored through the counter-electrode ion population, has relaxation time constants of only 1.2–3.5 ns, well below the corresponding DNA lie-down times (Fig. 3G,H: Supplementary Fig. S2). DNA lie-down can therefore be approximated as motion within an already established EDL under these conditions.

To characterize the EDL that forms ahead of DNA lie-down, we examine voltage-dependent ion and water distributions at MCH-passivated gold without DNA in 0.15 M NaCl (Fig. 3I–K). Negative and positive biases enrich Na^+^ and Cl^−^, respectively, near the outer edge of the MCH layer, while water layering outside the monolayer changes little. At stronger biases, however, ions and water penetrate the monolayer and reach the gold surface, with sodium penetration detected from −0.3 V and chloride penetration from +0.4 V. Such penetration may compromise passivation and device longevity, favoring operation at the lowest bias sufficient for switching. Where higher voltages are required, longer-chain SAMs should be considered to limit penetration.^42^

Across the voltage and salt series, lie-down is fastest at low salt and strong positive bias, whereas anionic MHA shifts the response toward higher voltages (Fig. 3L). At higher salt, increasing voltage does not produce a uniform acceleration: the response levels off at 0.15 M and becomes nonmonotonic at 1.0 M. We describe the switching fraction by *P*_switch_(*V*) = [1 + exp(*V* −(*V* – *V*_1_/_2_)/*w*)]^−1^, where *V*_1/2_ is the voltage at which 50% of the population switches and *b* describes the transition width. On MCH, *V*_1/2_ increases from approximately 0.06 to 0.10 and 0.28 V as NaCl increases from 0.01 to 0.15 and 1.0 M, respectively, whereas MHA reaches 0.40 V at 0.01 M (Fig. 3M). Low salt thus favors switching at lower voltage, while anionic passivation shifts the transition toward higher positive bias. This shift suggests a route to designing a semiconductor-like switching response with a tunable voltage threshold by controlling the charge density of the passivation layer. Note that these fitted voltages summarize unequal 50–600 ns observation windows and the prescribed surface ionization: they characterize switching under the sampled conditions rather than an intrinsic voltage threshold.

Although rotational orientations about the DNA helical axis upon surface contact may appear random across individual trajectories, statistical analysis of more than 200 lie-down trajectories reveals a strong preference. Across MCH and MHA under bias and AHT and bare gold without bias, the surface-facing direction peaks near the helical phases of bp10–bp11, with a smaller population spanning bp1–bp8 (Fig. 3N). Lie-down proceeds through a zip-like mechanism, in which contact begins near the anchor on the untethered strand and progresses toward the free end (Supplementary Fig. S3). During this process, dsDNA pivots about the anchor while rolling about its helical axis. After lie-down, the duplex lies along the surface (Supplementary Fig. S4). The preferred orientation offers a general structural basis for tethered DNA device design: for example, a redox reporter on the surface-facing side could gain electrode access more readily than one facing away, even at the same overall tilt.

### DNA recovery and surface-dependent release

Repeated electrical operation requires DNA recovery as well as lie-down. Removing the voltage bias allows DNA on MCH to stand up, but recovery slows markedly as salt concentration increases (Fig. 4A,B). Using sustained return to *θ* ≤ 75° to mark departure from the lying state, we find that the Kaplan–Meier median recovery time increases from 4.2 to 22.2 and 66.6 ns at 0.01, 0.15 and 1.0 M NaCl, respectively, an approximately sixteenfold increase. Unlike voltage-driven lie-down, this recovery is thermally driven once the bias is removed, yet remains strongly salt dependent.

**Fig. 4.**
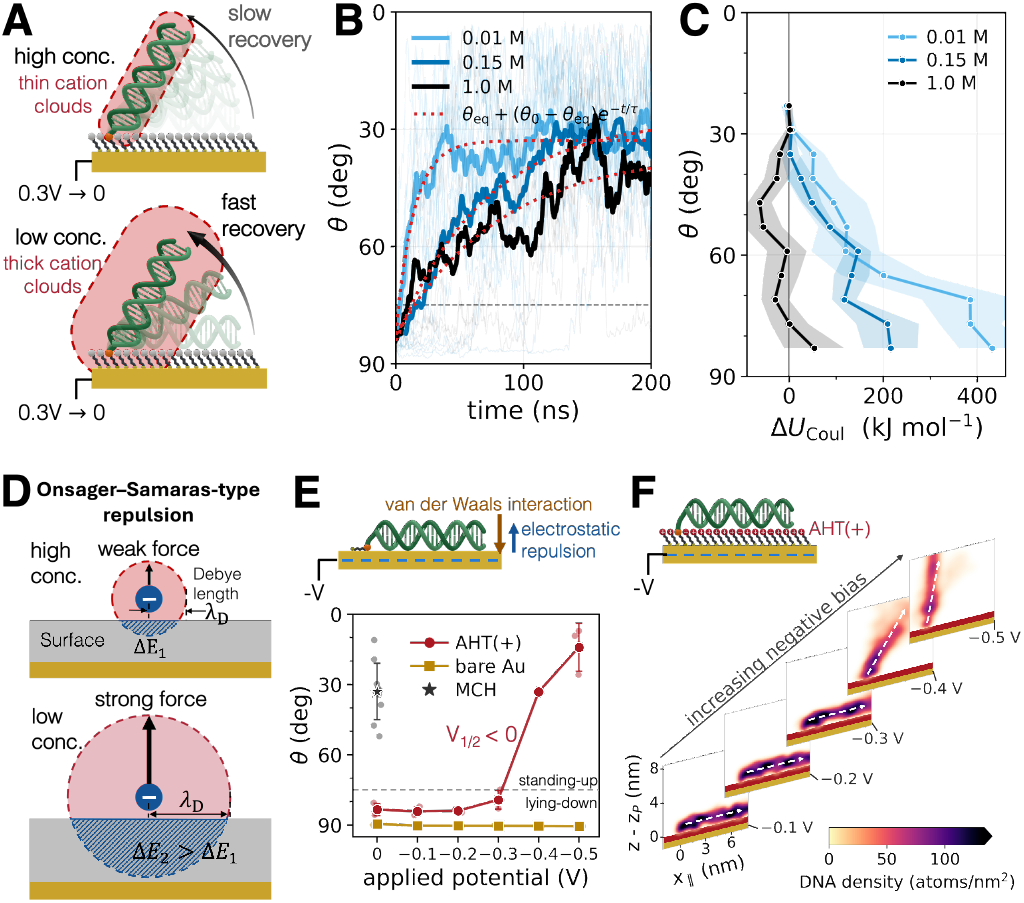
Interfacial repulsion and surface-dependent DNA recovery. **A**, Schematic of salt-dependent DNA recovery after bias removal. **B**, Salt-dependent recovery on MCH after switching from +0.3 to 0 V. Bold and faint traces denote ensemble means and individual trajectories: red dotted curves are guides to the eye. **C**, Short-range DNA–environment Coulomb energy on MCH at zero bias versus tilt, relative to the upright reference at *θ* = 20–32°. The environment includes water, ions and MCH. Points denote means across replicates: shading shows uncertainty estimates (Methods, “Contact face and Coulomb energy”). **D**, Schematic of counterion exclusion and its salt dependence. **E**, Mean tilt under negative bias on cationic AHT and bare gold at 0.1 M NaCl, with MCH at zero bias and 0.15 M as a reference. Faint points denote replicate means: error bars denote SD. The *V*_1/2_ *<* 0 annotation is qualitative. **F**, DNA density maps on AHT under negative bias at 0.1 M NaCl, using the molecular frame in Fig. 2C. White dashed arrows indicate dsDNA orientations. Dashed lines in B and E mark departure from the lying state, *θ* ≤ 75°. All simulations are at 310 K.

To probe the origin of this salt dependence, we analyzed the short-range DNA–environment Coulomb energy and found that the energy difference between lying configurations and the upright reference increases as salt concentration decreases (Fig. 4C). This salt-dependent energetic penalty reflects the same Onsager–Samaras-type repulsion that weakens surface-contact stabilization in the equilibrium free-energy analysis.^39^ As DNA approaches the surface, the space available to its surrounding counterions becomes restricted, weakening the screening of DNA charge and increasing the electrostatic cost of contact (Fig. 4D).^43,44^ This repulsion competes with attractive interactions between DNA and the terminal hydroxyl groups of MCH. At low salt, the counterion atmosphere extends farther, and its restriction near the surface carries a greater energetic cost. The resulting reduction in contact stability contributes to faster recovery and is consistent with the shallower near-contact free-energy minimum at 0.01 than at 0.15 M NaCl at zero bias (Fig. 2I).

The positively charged AHT monolayer and bare gold, in contrast, retain DNA near the surface at zero bias but respond differently to negative potentials (Fig. 4E,F). On AHT, pronounced recovery occurs around −0.4 V, comparable in magnitude and opposite in sign to the nominal +0.37 V equivalent of its positive surface charge. This correspondence suggests that opposing the cationic surface attraction enables release, giving rise to a negative switching voltage. As with MHA, this fixed 10% ionization means the switching voltage would similarly shift with pH and the interfacial *pK*_*a*_ in practice.^40,41^ DNA on bare gold, however, remains adsorbed even at −0.5 V. Short-range van der Waals attraction at multiple DNA–gold contacts can collectively stabilize adsorption against the opposing electrical force. For scale, single-molecule force spectroscopy of 30-nucleotide ssDNA homopolymers on aqueous Au(111) gives estimated adsorption free-energy magnitudes of approximately 21–36 kJ mol^−1^ per nucleotide, equivalent to the work on one elementary charge across 0.22–0.37 V.^45^ These experimental values indicate substantial binding at individual contacts: multiple contacts along dsDNA can collectively resist electrically driven release.

## Conclusions

Electrode-tethered dsDNA has been studied for nearly three decades, yet our ensemble all-atom simulations reveal molecular behavior and physical mechanisms that had gone unrecognized. For instance, the preferred orientation about the DNA helical axis upon surface contact was resolved through statistical analysis of more than two hundred trajectories in which DNA lies down on the surface. This result illustrates the value of an ensemble-level, all-atom approach. Onsager–Samaras-type repulsion, long established in electrolyte theory, further shows that DNA–surface interactions are not set by surface charge alone. Even on a net-neutral monolayer, restriction of the counterion atmosphere near the surface imposes an energetic cost on contact.

Taken together, these simulations reproduce the experimentally observed orientation and switching of tethered DNA and resolve the molecular interactions that underlie them. In doing so, they narrow the longstanding gap between molecular simulation and device-level measurement and point toward its closure. This molecular understanding of the interface provides a foundation for device design, connecting the salt concentration, surface chemistry and applied potential that experimentalists already control to the molecular interactions that shape contact, switching and release. Pursued alongside experiment, ensemble molecular dynamics could bring the same scrutiny to other well-studied tethered systems, advancing the molecular engineering of surface-tethered devices.

## Supporting information

Noh_TetheredDNA_SI

## Methods

### System construction

A 24 bp B-form duplex, 5*’* -TACCAGCTATTGTATCTAATAAGA-3*’* and its complement, was tethered at the 5*’* phosphate through a C6 thiolate linker to a three-layer Au(111) electrode, with a second Au(111) slab as the counter electrode. Surfaces were bare or coated with self-assembled monolayers of 6-mercapto-1-hexanol (MCH), 6-mercaptohexanoic acid (MHA) or 6-amino-1-hexanethiol (AHT). Monolayers contained 479 chains at 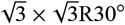 sites, with one central chain replaced by the linker: 10% of MHA and AHT chains were randomly assigned negative and positive formal charges, respectively. The 10.38×9.99×48.90 nm^3^ cell contained 4.2–4.6×10^4^ waters and a vacuum gap above the counter electrode. The vacuum spacing extended the box length along *z* to approximately three times the height of the electrode–electrolyte assembly. Together with the 3dc slab correction described below, this spacing suppresses spurious electrostatic interactions between periodic images along *z*. For DNA-containing systems, added NaCl concentrations were 0.01–1.0 M, calculated from water counts, with additional counterions for electroneutrality. Systems constructed under bias also contained additional NaCl pairs for double-layer formation: re-seeded ensembles retained their zero-bias parent composition (Supplementary Table S2). The labelled ensemble carried oxidized methylene blue (MB^+^) at the tethered strand’s 3*’* end through an alkyl linker.

### Force field

CHARMM36^50^ was used for DNA and CGenFF^51^ for monolayers, linkers and MB. Water was represented by TIP3P,^52^ with TIP4P/2005^53^ used for one comparison. Na^+^, Cl^−^ and gold used the July 2022 CHARMM36 parameters and pair-specific corrections: the AU Lennard-Jones parameters derive from Heinz et al.^54^ Gold was uncharged at zero bias: image-charge polarization^55^ was omitted. Each thiolate sulfur bonded to two bridge-site gold atoms using the parameters of Zhou et al.^56^ Gold atoms were restrained to their reference positions by harmonic potentials with a force constant of 20 000 kJ mol^−1^ nm^−2^ in each Cartesian direction: monolayer chains were unrestrained.

### Simulation protocol

Simulations used GPU-accelerated GROMACS 2025^57^ in the canonical ensemble at 310 K, with a velocity-rescaling thermostat^58^ (*τ*_*T*_ = 0.1 ps). Particle mesh Ewald^59^ used 0.15 nm Fourier spacing, the 3dc slab correction^60^ and *ϵ*_surface_ = 0. Real-space Coulomb and Lennard-Jones cutoffs were 1.2 nm. Lennard-Jones interactions used either a 1.0–1.2 nm force switch without dispersion correction or a potential shift with energy and pressure dispersion corrections, as specified by condition in Supplementary Table S1. A 4 fs timestep was used with threefold hydrogen mass repartitioning^61^ and LINCS constraints on bonds involving hydrogen. Steepest-descent minimization was followed by 2 ns relaxation at 310 K with uncharged electrodes and DNA position restraints of 100 kJ mol^−1^ nm^−2^. DNA restraints were then removed and the prescribed electrode charges applied. Replicate counts, trajectory durations and simulation conditions are listed in Supplementary Table S1.

### Applied potential

Applied bias was represented by equal and opposite fixed charges on the bottom working electrode and the top counter electrode, so that charging the electrodes introduced no net charge into the simulation cell. The working-electrode charge density was assigned as *σ*- = *C*_A_ *V*, using *C*_A_ = 20 *µ*F cm^−2^, selected at the lower end of the 20–30 *µ*F cm^−2^ range reported for bare Au(111) in aqueous Tris buffer.^62^ This value served as a common reference for converting imposed charge densities to nominal voltages across conditions, while the actual interfacial capacitance may vary with surface chemistry, electrolyte and potential. Positive *V* therefore gave positive charge on the bottom electrode and negative charge on the top electrode: reversing *V* reversed both signs. To approximate finite electronic screening within gold, the total charge on each slab was distributed across its three atomic layers using normalized Thomas–Fermi weights proportional to exp (−*d*_*i*_, /*λ*_TF_), where *d*_*i*_ is the depth of layer *i* from the electrolyte-facing surface and, *λ*_TF_ = 0.05 nm, a representative screening length for gold.^63^ This profile concentrates charge in the surface layer, with exponentially smaller contributions in deeper layers.^64^ Atomic charges remained fixed during each trajectory. Thus, *V* denotes the nominal bias corresponding to the imposed charge, rather than a calibrated or dynamically maintained electrode potential.^65,66^

### Orientation, switching and recovery

Trajectories were made whole across periodic boundaries. Tilt *θ* is the angle between the surface normal and the vector from the linker phosphate to the distal base-pair center. Equilibrium statistics used the last 100 ns, or the last half of shorter trajectories. Adsorbed distributions on AHT and bare gold included only trajectories reaching *θ* ≥ 70° for at least 5 ns, using frames after that event. Driven lie-down and recovery were the first crossings of *θ* ≥ 75° and *θ* ≤ 75°, respectively, sustained for 5 ns after a 1 ns moving average. Trajectories without events were right-censored at their final time. Lie-down comparisons use Kaplan–Meier medians in Fig. 3L and observed-event medians in Fig. 3H: recovery times are Kaplan–Meier medians (Supplementary Tables S3 and S4).

The fractions of trajectories with lie-down events within their observation windows were fitted by binomial maximum likelihood, with *w* ≥ 0.02 V and 95% likelihood intervals for *V*_1/2_ at the fitted width. The zero-voltage point used the equilibrium ensemble, including 0.1 M for the 0.15 M series. Double-layer relaxation was fitted exponentially to the ensemble-mean sodium population at 12 ≤ *z* ≤ 15 nm near the counter electrode: intervals used the 16th–84th percentiles of 1000 trajectory bootstrap resamples. Unless specified in the captions, uncertainties denote SEM across independent replicates.

### Structure, fraying and apparent moduli

Base-pair frames and local step parameters followed standard geometric definitions.^67,68^ End-to-end length was measured between basepair-frame origins 1 and 24, with a B-form reference of 7.82 nm. Length and mean twist used frames after 20 ns, with twist averaged over steps 5–20: adjacent-origin spacing used all frames and local twist used locally paired steps. A base pair was open when at least two thirds of its Watson–Crick donor–acceptor distances exceeded 3.5 Å. Terminal fraying was quantified by counting consecutive open pairs inward from each terminus over the last 150 ns.

Moduli were estimated by inverse-covariance analysis of conformational fluctuations,^69^ using 11–14-step fragments sampled within bp 5–21 or bp 1–24, with identical frames and fragment lengths for both windows. Canonical frames required no open pair over bp 5– 21 and local twist above 10° at every step from 5 to 20. Replicates required at least 500 canonical frames, of which at least 50 fell within the last 150 ns used for analysis. Fluctuations were centered on each fragment’s mean before pooling. For an N-step fragment, *L*_0_ = 0.34*N* nm, *S* = *k*_B_*T L*_0_/Var_*p*_ (*L*), *C* = *k*_B_*T L*_0_/Var_*p*_ (Φ) and *B* = *k*_B_*T L*_0_/{[Var_*p*_ (roll) + Var_*p*_ (tilt)]/2}, where Φ is end-to-end length, Φ is summed helical twist, and roll and tilt describe the relative orientation of the fragment’s terminal base-pair frames: all angles are in radians. Conditional variances were Var_*p*_(*x*_*i*_) = 1/(∑^−1^) for the respective covariance matrix Σ: stretch was conditioned on summed helical twist and helical-axis bend, torsion on summed helical rise and helical-axis bend, and roll and tilt on each other, summed helical twist and contour length. Estimates were averaged over fragment lengths within each replicate, then across replicates.

### Free-energy landscape and joint geometry

Simulations at different electrode potentials were combined to improve sampling of tilt angles rarely visited at zero bias. The voltage-dependent free-energy landscape was represented as *F* (*θ, V*) = *F*_0_ (*θ*)+ *A W* (*θ, V*), where *F*_0_ is the zero-bias landscape and *A W* describes the electrostatic contribution of the applied bias. This contribution was calculated using the nonlinear Gouy–Chapman potential,^70^

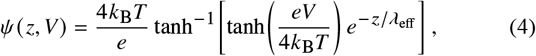

where *z* is the distance from the model interface, *e* is the elementary charge, and the nominal bias *V* serves as the model boundary potential relative to the bulk. The effective screening length was 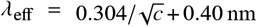, where *c* is the added NaCl concentration in molar units and the 0.40 nm correction accounts for hydration. For a rigid rod of length *L* = 8.16 nm tilted by *θ* from the surface normal,

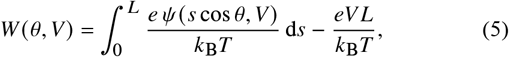

with the reference chosen such that *W* (90°, *V*) = 0. Angular distributions were fitted jointly by multinomial maximum likelihood71 to determine *F*0 and a salt-specific coupling coefficient *A* incorporating the effective DNA charge density, with *A W* expressed in energy units.

Histograms used 2° bins, with frames from 200–1000 ns at 0.15 M and −0.3 to +0.3 V. At 0.01 and 1.0 M, zero-bias ensembles were combined with the final 100 ns of voltage-step trajectories. Quantitative analysis at 1.0 M was restricted to *θ* ≤ 70°.

The orientational contribution was removed through *U* (*θ*) = *F*_0_ (*θ*)+ *k*B*T* ln sin *θ*. At 0.15 M, *U* was fitted over 0–70° to the hyperboliccosine potential in Eq. 2, yielding *a* = 1.905 and *b* = 0.732 rad. The residual *U* − *U*tilt defined the effective DNA–surface contribution.

Joint geometry was evaluated over 20–200 ns from five zero-bias MCH replicates at 0.1 M. The duplex axis was defined by the principal axis of the 24 base-pair C1*’* centers, directed away from the tether. The joint angle *φ* was measured between this axis and the linker sulfur-to-phosphate vector. Signed bending was defined as the polar tilt of the distal axis (bp 13–24) minus that of the proximal axis (bp 1–12). The effective joint potential, − *k*B*T* ln[*P*(*φ*)/sin *φ*], was calculated in 2.5° bins for each replicate and then averaged. In Fig. 2J, bending error bars are the population SD of pooled angles within each 5° tilt bin divided by 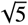. Joint-angle and joint-potential error bars (Fig. 2J and L) are the population SD of per-replicate bin means and potentials, respectively, divided by 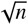, where *n* is the number of contributing replicates in each bin.

### Contact face and Coulomb energy

Contact-face analysis used the last 10 ns of trajectories with *θ* > 60°. The downward surface normal and cross-strand vectors were projected perpendicular to an axis fitted to paired C1*’* centers at bp 8–23: their signed angles were corrected for the mean helical winding phase of each base pair. A base pair counted as in contact when its C1*’* separation was 0.8–1.4 nm and its lowest heavy atom lay within 0.8 nm of the contact floor, the second percentile of per-frame minimum base-pair heights for that condition. Trajectories with at least ten contact frames contributed equally to the circular mean.

DNA–environment Coulomb energies included short-range interactions with water, Na^+^, Cl^−^ and MCH, excluding long-range PME. Energies sampled every 100 ps were averaged in 6° tilt bins after subtracting each replicate’s mean over 20–32°. Bin means used contributing replicates: shaded half-widths were their sample SD divided by 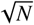, with total retained ensemble sizes *N* = 10, 10 and 5 at 0.01, 0.15 and 1.0 M, respectively.

## Data availability

The data supporting the findings of this study are available from the corresponding authors upon reasonable request.

## Acknowledgements

Y.N. acknowledges support from the University of Notre Dame Bio-engineering and Life Sciences (BELS) Postdoctoral Fellowship.

## Author contributions

Y.N.: Conceptualization, Methodology, Software, Investigation, Formal analysis, Data curation, Visualization, Funding acquisition, Resources, Project administration, Writing – original draft, Writing – review & editing.

H.C.: Resources, Project administration, Writing – review & editing. K.F.: Conceptualization, Resources, Project administration, Supervision, Writing – review & editing.

