## Supplementary material for "Electrode-Tethered DNA: Interfacial Mechanics and Switching Dynamics from Ensemble-Level Atomistic Simulations": Noh_TetheredDNA_SI

### Supporting information

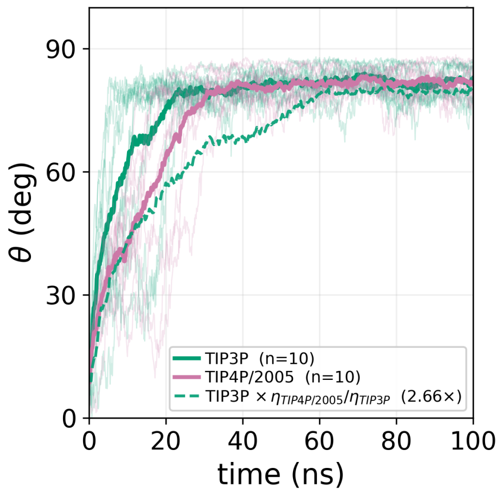


**Supplementary Fig. S1. Water-model dependence of DNA lie-down.** DNA on MCH at nominal +0.3 V, 0.01 M NaCl and 310 K. Thin and thick curves denote individual trajectories ($n = 10$ per water model) and their means, respectively. The tilt angle $\theta$ is measured from the surface normal, as in the main text. The dashed curve multiplies TIP3P times by 2.66, the TIP4P/2005-to-TIP3P bulk viscosity ratio at 298 K and 1 bar,[72] for a qualitative comparison.

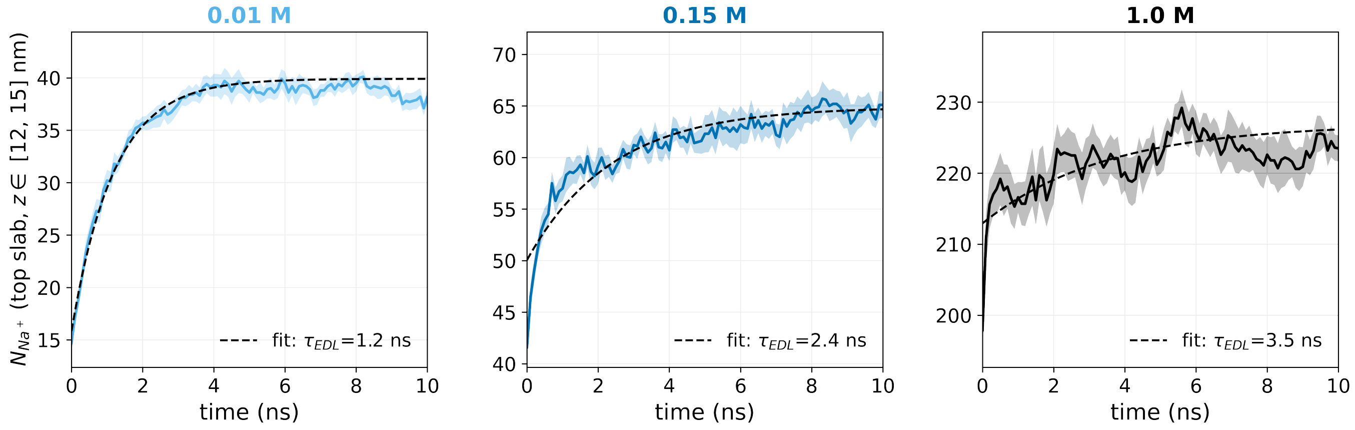


**Supplementary Fig. S2. Counterelectrode double-layer relaxation.** Sodium population within $12 \leq z \leq 15$ nm after application of nominal +0.3 V at 0.01, 0.15 and 1.0 M NaCl and 310 K. Solid curves and shading denote mean ± SEM across ten trajectories. Dashed curves are exponential fits with $\tau_{\mathrm{EDL}} = 1.2$, 2.4 and 3.5 ns, respectively. Fits use the full 100, 200 and 100 ns trajectories; the first 10 ns are shown.

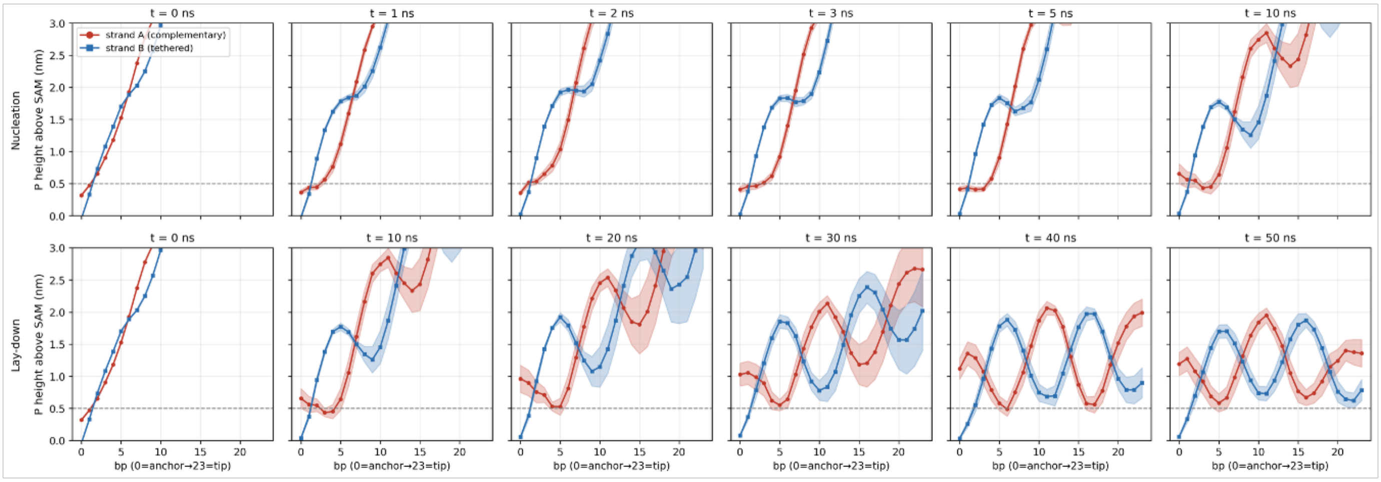


**Supplementary Fig. S3. Strand-resolved height profiles during zip-like lie-down.** DNA on MCH at nominal +0.2 V and 0.01 M NaCl. Curves and shading denote mean ± SEM across ten trajectories for the complementary strand A (red) and tethered strand B (blue). Phosphate heights are measured above the SAM, with terminal O5′ atoms used where phosphate atoms are absent. Base-pair indices run from the anchor (0) to the free end (23). The upper and lower rows cover 0–10 and 0–50 ns, respectively; times are measured from the start of the trajectories. Dashed lines mark a height of 0.5 nm.

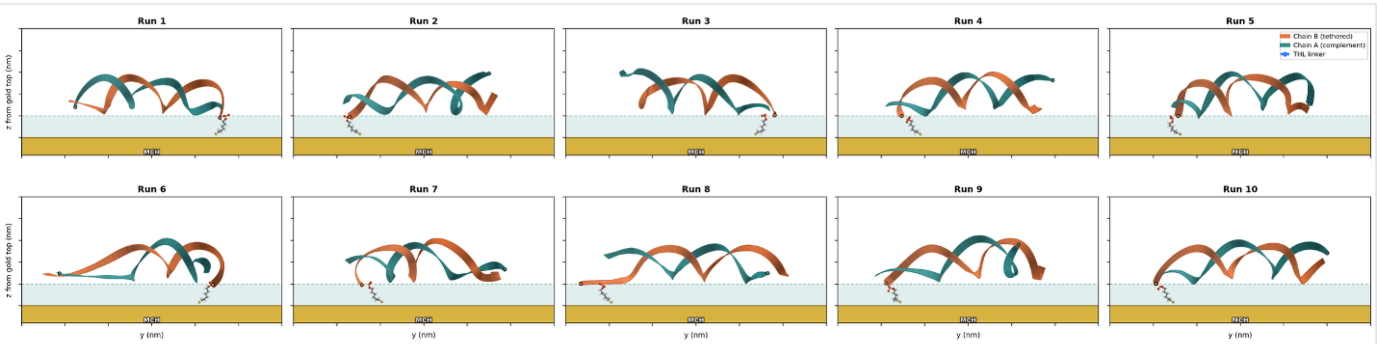


**Supplementary Fig. S4. DNA configurations after lie-down across ten trajectories.** Side views at 100 ns on MCH at nominal +0.3 V and 0.01 M NaCl. Orange and teal ribbons denote tethered strand B and complementary strand A, respectively; the thiol linker is shown explicitly. Each panel corresponds to one trajectory.

### Supporting tables

#### Simulation conditions

Supplementary Table S1 includes all 620 trajectories across 187 condition labels used to develop this study, including initial system and protocol tests and electrode controls without DNA. The total sampled time is approximately 144 $\mu$s after joining continuations and counting overlapping intervals and duplicate copies once. Entries without a log or trajectory confirming execution are excluded.

Target durations are the planned lengths per trajectory, including extensions; individual early termination times are omitted. The total sampled time above uses the actual completed intervals. PS denotes a Lennard-Jones potential shift with energy and pressure dispersion corrections; FS denotes a force switch without dispersion correction. A dash denotes an unavailable entry. Condition IDs follow the source inventory. 1MH denotes 1-mercaptohexane. Recorded water models are TIP3P except C161 (TIP4P/2005); the water model is unspecified in the early setup records C172, C173, C202, C203, C207, C232 and C241. Each re-seeded ensemble contains 50 trajectories started from ten configurations of each of five parent trajectories.

Supplementary Table S1: Complete simulation inventory. $n$ is the number of trajectories after joining continuations; target durations are given as number of trajectories × planned length, with units indicated. NaCl concentrations and electrode biases are nominal.

| ID | System | NaCl (M) | Bias (V) | $n$ | Target duration | Tail |
|---|---|---|---|---|---|---|
| *Initial DNA and linker tests* | | | | | | |
| C172 | DNA in water | – | – | 1 | 1×50 ps | – |
| C173 | Thiol-linked DNA in water | – | – | 1 | 1×50 ps | – |
| *Initial gold–DNA tests* | | | | | | |
| C202 | Au–DNA, setup test | – | – | 1 | 1×5 ps | – |
| C203 | Au–DNA, fixed-gold test | – | – | 1 | 1×10 ps | – |
| *Reflective-wall test* | | | | | | |
| C207 | Au–DNA, reflective wall | – | – | 1 | 1×5 ps | – |
| *GolP test* | | | | | | |
| C232 | Au–DNA, GolP test | – | – | 1 | 1×2 ps | – |
| *Initial SAM–DNA test* | | | | | | |
| C241 | SAM–DNA, setup test | – | – | 1 | 1×5 ps | – |
| *MHA charge series* | | | | | | |
| C001 | MHA, neutral | – | 0 | 2 | 2×500 ns | PS |
| C003 | MHA, 30.1% charged | – | 0 | 2 | 2×500 ns | PS |
| C005 | MHA, 20.0% charged | – | 0 | 2 | 2×500 ns | PS |
| C007 | MHA, 10.0% charged | – | 0 | 2 | 2×500 ns | PS |
| *AHT charge series and initial electrode controls* | | | | | | |
| C010 | AHT, neutral | – | 0 | 2 | 2×500 ns | PS |
| C012 | AHT, 30.1% charged | – | 0 | 2 | 2×500 ns | PS |
| C014 | AHT, 20.0% charged | – | 0 | 2 | 2×500 ns | PS |
| C016 | AHT, 10.0% charged | – | 0 | 2 | 2×500 ns | PS |
| C018 | AHT, 5.0% charged | – | 0 | 2 | 2×500 ns | PS |
| C020 | AHT, 2.5% charged | – | 0 | 2 | 2×500 ns | PS |
| C021 | MCH, no DNA | – | 0 | 1 | 1×20 ns | PS |
| C022 | MCH, no DNA | – | 0.1 | 1 | 1×20 ns | PS |
| C023 | MCH, no DNA | – | 0.2 | 1 | 1×20 ns | PS |
| C024 | MCH, no DNA | – | 0.3 | 1 | 1×20 ns | PS |
| C025 | MCH, no DNA | – | 0.4 | 1 | 1×20 ns | PS |
| C026 | MCH, no DNA | – | 0.5 | 1 | 1×20 ns | PS |
| C027 | MCH, no DNA | – | 0.6 | 1 | 1×20 ns | PS |
| C028 | MCH, no DNA | – | 0.7 | 1 | 1×20 ns | PS |
| C029 | MCH, no DNA | – | 0.8 | 1 | 1×20 ns | PS |
| C030 | MCH, no DNA | – | −0.1 | 1 | 1×20 ns | PS |
| C031 | MCH, no DNA | – | −0.2 | 1 | 1×20 ns | PS |
| C032 | MCH, no DNA | – | −0.3 | 1 | 1×20 ns | PS |
| C033 | MCH, no DNA | – | −0.4 | 1 | 1×20 ns | PS |
| C034 | MCH, no DNA | – | −0.5 | 1 | 1×20 ns | PS |
| C035 | MCH, no DNA | – | −0.6 | 1 | 1×20 ns | PS |
| C036 | MCH, no DNA | – | −0.7 | 1 | 1×20 ns | PS |
| C037 | MCH, no DNA | – | −0.8 | 1 | 1×20 ns | PS |
| *Initial MCH switching series* | | | | | | |
| C038 | MCH | – | 0.4 | 1 | 1×400 ns | – |
| C039 | MCH | – | 0.6 | 1 | 1×200 ns | – |
| C040 | MCH | – | 0.7 | 1 | 1×50 ns | – |
| C041 | MCH | – | 0.8 | 1 | 1×50 ns | – |
| *Electrode controls without DNA* | | | | | | |
| C042 | MCH, no DNA | 0.015 | 0.3 | 1 | 1×100 ns | PS |
| C043 | MCH, no DNA | 0.05 | 0.3 | 1 | 1×100 ns | PS |
| C044 | MCH, no DNA | 0.15 | 0.3 | 1 | 1×100 ns | PS |
| C045 | MCH, no DNA | 0.3 | 0.3 | 1 | 1×100 ns | PS |
| C046 | MCH, no DNA | 1 | 0.3 | 1 | 1×100 ns | PS |
| C047 | MCH, no DNA | 0.15 | 0 | 1 | 1×100 ns | PS |
| C048 | MCH, no DNA | 0.15 | 0.1 | 1 | 1×100 ns | PS |
| C049 | MCH, no DNA | 0.15 | 0.2 | 1 | 1×100 ns | PS |
| C050 | MCH, no DNA | 0.15 | 0.3 | 1 | 1×100 ns | PS |
| C051 | MCH, no DNA | 0.15 | 0.4 | 1 | 1×100 ns | PS |
| C052 | MCH, no DNA | 0.15 | 0.5 | 1 | 1×100 ns | PS |
| C053 | MCH, no DNA | 0.15 | −0.1 | 1 | 1×100 ns | PS |
| C054 | MCH, no DNA | 0.15 | −0.2 | 1 | 1×100 ns | PS |

Supplementary Table S1 (continued)

| ID | System | NaCl (M) | Bias (V) | $n$ | Target duration | Tail |
|---|---|---|---|---|---|---|
| C055 | MCH, no DNA | 0.15 | −0.3 | 1 | 1×100 ns | PS |
| C056 | MCH, no DNA | 0.15 | −0.4 | 1 | 1×100 ns | PS |
| C057 | MCH, no DNA | 0.15 | −0.5 | 1 | 1×100 ns | PS |
| C058 | bare Au, no DNA | 0.15 | 0 | 1 | 1×100 ns | PS |
| C059 | bare Au, no DNA | 0.15 | 0.1 | 1 | 1×100 ns | PS |
| C060 | bare Au, no DNA | 0.15 | 0.2 | 1 | 1×100 ns | PS |
| C061 | bare Au, no DNA | 0.15 | 0.3 | 1 | 1×100 ns | PS |
| C062 | bare Au, no DNA | 0.15 | 0.4 | 1 | 1×100 ns | PS |
| C063 | bare Au, no DNA | 0.15 | 0.5 | 1 | 1×100 ns | PS |
| C064 | bare Au, no DNA | 0.15 | −0.1 | 1 | 1×100 ns | PS |
| C065 | bare Au, no DNA | 0.15 | −0.2 | 1 | 1×100 ns | PS |
| C066 | bare Au, no DNA | 0.15 | −0.3 | 1 | 1×100 ns | PS |
| C067 | bare Au, no DNA | 0.15 | −0.4 | 1 | 1×100 ns | PS |
| C068 | bare Au, no DNA | 0.15 | −0.5 | 1 | 1×100 ns | PS |
| C069 | MCH, no DNA | – | 0 | 1 | 1×20 ns | PS |
| C070 | MCH, no DNA | – | 0.1 | 1 | 1×20 ns | PS |
| C071 | MCH, no DNA | – | 0.2 | 1 | 1×20 ns | PS |
| C072 | MCH, no DNA | – | 0.3 | 1 | 1×20 ns | PS |
| C073 | MCH, no DNA | – | 0.4 | 1 | 1×20 ns | PS |
| C074 | MCH, no DNA | – | 0.5 | 1 | 1×20 ns | PS |
| C075 | MCH, no DNA | – | 0.6 | 1 | 1×20 ns | PS |
| C076 | MCH, no DNA | – | 0.7 | 1 | 1×20 ns | PS |
| C077 | MCH, no DNA | – | 0.8 | 1 | 1×20 ns | PS |
| C078 | MCH, no DNA | – | −0.1 | 1 | 1×20 ns | PS |
| C079 | MCH, no DNA | – | −0.2 | 1 | 1×20 ns | PS |
| C080 | MCH, no DNA | – | −0.3 | 1 | 1×20 ns | PS |
| C081 | MCH, no DNA | – | −0.4 | 1 | 1×20 ns | PS |
| C082 | MCH, no DNA | – | −0.5 | 1 | 1×20 ns | PS |
| C083 | MCH, no DNA | – | −0.6 | 1 | 1×20 ns | PS |
| C084 | MCH, no DNA | – | −0.7 | 1 | 1×20 ns | PS |
| C085 | MCH, no DNA | – | −0.8 | 1 | 1×20 ns | PS |
| C086 | MCH, no DNA | 0.15 | 0 | 1 | 1×40 ns | PS |
| C087 | MCH, no DNA | 0.15 | 0.1 | 1 | 1×40 ns | PS |
| C088 | MCH, no DNA | 0.15 | 0.2 | 1 | 1×40 ns | PS |
| C089 | MCH, no DNA | 0.15 | 0.3 | 1 | 1×40 ns | PS |
| C090 | MCH, no DNA | 0.15 | 0.4 | 1 | 1×40 ns | PS |
| C091 | MCH, no DNA | 0.15 | 0.5 | 1 | 1×40 ns | PS |
| C092 | MCH, no DNA | 0.15 | −0.1 | 1 | 1×40 ns | PS |
| C093 | MCH, no DNA | 0.15 | −0.2 | 1 | 1×40 ns | PS |
| C094 | MCH, no DNA | 0.15 | −0.3 | 1 | 1×40 ns | PS |
| C095 | MCH, no DNA | 0.15 | −0.4 | 1 | 1×40 ns | PS |
| C096 | MCH, no DNA | 0.15 | −0.5 | 1 | 1×40 ns | PS |
| C097 | bare Au, no DNA | 0.15 | 0 | 1 | 1×40 ns | PS |
| C098 | bare Au, no DNA | 0.15 | 0.1 | 1 | 1×40 ns | PS |
| C099 | bare Au, no DNA | 0.15 | 0.2 | 1 | 1×40 ns | PS |
| C100 | bare Au, no DNA | 0.15 | 0.3 | 1 | 1×40 ns | PS |
| C101 | bare Au, no DNA | 0.15 | 0.4 | 1 | 1×40 ns | PS |
| C102 | bare Au, no DNA | 0.15 | 0.5 | 1 | 1×40 ns | PS |
| C103 | bare Au, no DNA | 0.15 | −0.1 | 1 | 1×40 ns | PS |
| C104 | bare Au, no DNA | 0.15 | −0.2 | 1 | 1×40 ns | PS |
| C105 | bare Au, no DNA | 0.15 | −0.3 | 1 | 1×40 ns | PS |
| C106 | bare Au, no DNA | 0.15 | −0.4 | 1 | 1×40 ns | PS |
| C107 | bare Au, no DNA | 0.15 | −0.5 | 1 | 1×40 ns | PS |
| *Salt and ion-penetration controls without DNA* | | | | | | |
| C108 | MCH, no DNA | 0.05 | 0 | 1 | 1×100 ns | PS |
| C109 | MCH, no DNA | 0.15 | 0 | 1 | 1×100 ns | PS |
| C110 | MCH, no DNA | 0.3 | 0 | 1 | 1×100 ns | PS |
| C111 | MCH, no DNA | 1 | 0 | 1 | 1×100 ns | PS |
| C113 | MCH, no DNA | 0.15 | 0.5 | 1 | 1×200 ns | PS |
| C114 | MCH, no DNA | 0.3 | 0.5 | 1 | 1×200 ns | PS |
| C115 | MCH, no DNA | 0.5 | 0.5 | 1 | 1×200 ns | PS |
| C116 | MCH, no DNA | 1 | 0.5 | 1 | 1×200 ns | PS |
| C117 | MCH, no DNA | 1.5 | 0.5 | 1 | 1×200 ns | PS |
| *MCH voltage sweep* | | | | | | |
| C118 | MCH | 0.15 | 0 | 3 | 3×1000 ns | PS |
| C119 | MCH | 0.15 | 0.1 | 2 | 2×1000 ns | PS |
| C120 | MCH | 0.15 | 0.2 | 2 | 2×1000 ns | PS |
| C121 | MCH | 0.15 | 0.3 | 2 | 2×1000 ns | PS |
| C122 | MCH | 0.15 | 0.4 | 2 | 2×1000 ns | PS |
| C123 | MCH | 0.15 | 0.5 | 2 | 2×1000 ns | PS |
| C124 | MCH | 0.15 | −0.1 | 2 | 2×1000 ns | PS |
| C125 | MCH | 0.15 | −0.2 | 2 | 2×1000 ns | PS |
| C126 | MCH | 0.15 | −0.3 | 2 | 2×1000 ns | PS |
| *Bare-gold voltage and salt series* | | | | | | |
| C127 | bare Au | 0.01 | 0 | 5 | 4×400 ns, 1×200 ns | FS |
| C128 | bare Au | 0.15 | 0 | 2 | 2×1000 ns | PS |
| C129 | bare Au | 0.1 | 0 | 5 | 3×400 ns, 2×200 ns | FS |
| C130 | bare Au | 1.0 | 0 | 5 | 2×400 ns, 3×200 ns | FS |
| C131 | bare Au | 0.15 | 0.1 | 2 | 2×1000 ns | PS |
| C132 | bare Au | 0.15 | 0.2 | 2 | 2×1000 ns | PS |
| C133 | bare Au | 0.15 | 0.3 | 2 | 2×1000 ns | PS |
| C134 | bare Au | 0.15 | 0.4 | 2 | 2×1000 ns | PS |
| C135 | bare Au | 0.15 | 0.5 | 2 | 2×1000 ns | PS |

Supplementary Table S1 (continued)

| ID | System | NaCl (M) | Bias (V) | $n$ | Target duration | Tail |
|---|---|---|---|---|---|---|
| C136 | bare Au | 0.15 | −0.1 | 2 | 2×1000 ns | PS |
| C137 | bare Au | 0.15 | −0.2 | 2 | 2×1000 ns | PS |
| C138 | bare Au | 0.15 | −0.3 | 2 | 2×1000 ns | PS |
| *MCH switching, recovery and contact ensembles* | | | | | | |
| C139 | AHT(+) | 0.1 | −0.1 | 3 | 3×200 ns | FS |
| C140 | AHT(+) | 0.1 | −0.2 | 3 | 3×200 ns | FS |
| C141 | AHT(+) | 0.1 | −0.3 | 3 | 3×200 ns | FS |
| C142 | AHT(+) | 0.1 | −0.4 | 3 | 3×200 ns | FS |
| C143 | AHT(+) | 0.1 | −0.5 | 3 | 3×200 ns | FS |
| C144 | MCH | 0.01 | 0.1 | 10 | 8×400 ns, 2×200 ns | PS |
| C145 | MCH | 1.0 | 0.1 | 10 | 10×400 ns | FS |
| C146 | MCH | 1.0 | 0.2 | 10 | 10×400 ns | PS |
| C147 | MCH | 1.0 | 0.3 | 10 | 10×400 ns | PS |
| C148 | MCH | 0.15 | 0.1 | 10 | 10×400 ns | FS |
| C149 | MCH | 0.15 | 0.2 | 10 | 4×400 ns, 6×200 ns | PS |
| C150 | MCH | 1.0 | 0.4 | 10 | 10×400 ns | FS |
| C151 | MCH | 0.15 | 0.5 | 10 | 4×400 ns, 6×50 ns | PS |
| C152 | MCH | 1.0 | 0.5 | 10 | 9×400 ns, 1×100 ns | PS |
| C153 | bare Au | 0.1 | −0.1 | 3 | 3×200 ns | FS |
| C154 | bare Au | 0.1 | −0.2 | 3 | 3×200 ns | FS |
| C155 | bare Au | 0.1 | −0.3 | 3 | 3×200 ns | FS |
| C156 | bare Au | 0.1 | −0.4 | 3 | 3×200 ns | FS |
| C157 | bare Au | 0.1 | −0.5 | 3 | 3×200 ns | FS |
| C158 | MCH | 0.01 | 0.2 | 10 | 10×100 ns | PS |
| C159 | MCH, re-seeded | 0.01 | 0.2 | 50 | 50×50 ns | FS |
| C160 | MCH | 0.01 | 0.3 | 10 | 10×100 ns | PS |
| C161 | MCH, TIP4P/2005 | 0.01 | 0.3 | 10 | 10×100 ns | PS |
| C162 | MCH | 0.15 | 0.3 | 10 | 10×200 ns | PS |
| C163 | MCH | 0.01 | 0.4 | 10 | 10×50 ns | PS |
| C164 | MCH | 0.15 | 0.4 | 10 | 10×100 ns | PS |
| C165 | MCH | 0.01 | 0.5 | 10 | 10×100 ns | PS |
| C166 | MCH | 0.01 | −0.1 | 5 | 5×200 ns | FS |
| C167 | MCH | 0.01 | −0.2 | 5 | 5×200 ns | FS |
| C168 | MCH | 0.01 | −0.3 | 5 | 5×200 ns | FS |
| C169 | MCH | 0.01 | 0 after +0.3 | 10 | 10×200 ns | PS |
| C170 | MCH | 0.15 | 0 after +0.3 | 10 | 10×200 ns | PS |
| C171 | MCH | 1.0 | 0 after +0.3 | 5 | 5×200 ns | PS |
| *Zero-bias surface and salt series* | | | | | | |
| C174 | 1MH | 0.01 | 0 | 5 | 5×200 ns | FS |
| C175 | 1MH | 0.1 | 0 | 5 | 5×200 ns | FS |
| C176 | 1MH | 1.0 | 0 | 5 | 5×200 ns | FS |
| C177 | AHT(+) | 0.01 | 0 | 5 | 5×200 ns | FS |
| C178 | AHT(+) | 0.1 | 0 | 5 | 5×200 ns | FS |
| C179 | AHT(+) | 1.0 | 0 | 5 | 2×400 ns, 3×200 ns | FS |
| C180 | MCH | 0.01 | 0 | 5 | 5×200 ns | FS |
| C181 | MCH | 0.1 | 0 | 5 | 5×200 ns | FS |
| C182 | MCH | 1.0 | 0 | 5 | 5×200 ns | FS |
| C183 | MHA(−) | 0.01 | 0 | 5 | 5×200 ns | FS |
| C184 | MHA(−) | 0.1 | 0 | 5 | 5×200 ns | FS |
| C185 | MHA(−) | 1.0 | 0 | 5 | 5×200 ns | FS |
| *MHA switching series* | | | | | | |
| C186 | MHA(−) | 0.01 | 0.2 | 10 | 10×300 ns | FS |
| C187 | MHA(−) | 0.01 | 0.3 | 10 | 10×300 ns | FS |
| C188 | MHA(−) | 0.01 | 0.4 | 10 | 6×600 ns, 4×300 ns | FS |
| C189 | MHA(−) | 0.01 | 0.5 | 10 | 1×600 ns, 9×300 ns | FS |
| *Methylene-blue equilibrium series* | | | | | | |
| C190 | MCH+MB | 0.01 | 0 | 5 | 5×100 ns | FS |
| C191 | MCH+MB | 0.1 | 0 | 5 | 5×200 ns | FS |
| C192 | MCH+MB | 1.0 | 0 | 5 | 5×200 ns | FS |
| *Methylene-blue contact ensemble* | | | | | | |
| C193 | MCH+MB, re-seeded | 0.01 | 0.3 | 50 | 50×50 ns | FS |
| Total: 187 condition labels | | | | 620 | Sampled: 144 $\mu$s | |

#### Salt concentrations and ion counts

Solution ions include the added NaCl and the counterions required to neutralize DNA, charged monolayers and electrodes. We estimate volume-averaged concentrations as $c_s = N_s/(0.602214076V_w)$ M and $I = (c_{\mathrm{Na}} + c_{\mathrm{Cl}})/2$, using the water volume $V_w = N_w/33.4\ \mathrm{nm}^3$. Here $N_s$ and $N_w$ are the ion and water counts, respectively. Recovery and re-seeded trajectories retain the ion counts of their parent configurations.

Supplementary Table S2: Ion and water counts and estimated ionic strengths. Counts are identical across the $n$ configurations checked for ion composition in each condition.

| Surface | Label (M) | $V$ (V) | $n$ | $N_{\mathrm{Na}}$ | $N_{\mathrm{Cl}}$ | $N_w$ | $I$ (M) |
|---|---|---|---|---|---|---|---|
| Bare Au | 0.01 | 0.0 | 5 | 56 | 9 | 46485 | 0.0388 |
| Bare Au | 0.15 | 0.0 | 2 | 173 | 126 | 46251 | 0.1793 |
| Bare Au | 0.1 | 0.0 | 5 | 131 | 84 | 46335 | 0.1287 |
| Bare Au | 1.0 | 0.0 | 5 | 886 | 839 | 44825 | 1.0672 |
| AHT recovery | 0.1 | -0.4 | 3 | 79 | 80 | 43417 | 0.1016 |
| MCH | 1.0 | 0.1 | 10 | 841 | 794 | 42278 | 1.0724 |
| MCH | 0.15 | 0.1 | 10 | 169 | 122 | 43622 | 0.1850 |
| Bare Au recovery | 0.1 | -0.4 | 3 | 131 | 84 | 46335 | 0.1287 |
| MCH | 0.01 | 0.1 | 8 | 63 | 16 | 43834 | 0.0500 |
| MCH | 0.01 | 0.2 | 10 | 75 | 28 | 43810 | 0.0652 |
| MCH | 0.15 | 0.2 | 3 | 176 | 129 | 43608 | 0.1940 |
| MCH | 1.0 | 0.2 | 10 | 844 | 797 | 42272 | 1.0765 |
| MCH | 0.01 | 0.3 | 10 | 87 | 40 | 43786 | 0.0804 |
| MCH | 0.15 | 0.3 | 10 | 186 | 139 | 43588 | 0.2068 |
| MCH | 1.0 | 0.3 | 10 | 849 | 802 | 42262 | 1.0833 |
| MCH | 0.01 | 0.4 | 10 | 100 | 53 | 43760 | 0.0970 |
| MCH | 0.15 | 0.4 | 10 | 197 | 150 | 43566 | 0.2209 |
| MCH | 1.0 | 0.4 | 10 | 855 | 808 | 42250 | 1.0915 |
| MCH | 0.01 | 0.5 | 10 | 113 | 66 | 43734 | 0.1135 |
| MCH | 0.15 | 0.5 | 10 | 208 | 161 | 43544 | 0.2350 |
| MCH | 1.0 | 0.5 | 10 | 863 | 816 | 42234 | 1.1024 |
| MCH | 0.01 | -0.1 | 5 | 63 | 16 | 43834 | 0.0500 |
| MCH | 0.01 | -0.2 | 5 | 75 | 28 | 43810 | 0.0652 |
| MCH | 0.01 | -0.3 | 5 | 87 | 40 | 43786 | 0.0804 |
| AHT | 0.01 | 0.0 | 5 | 8 | 9 | 43559 | 0.0108 |
| AHT | 0.1 | 0.0 | 5 | 79 | 80 | 43417 | 0.1016 |
| AHT | 1.0 | 0.0 | 5 | 786 | 787 | 42003 | 1.0385 |
| MCH | 0.01 | 0.0 | 5 | 55 | 8 | 43850 | 0.0398 |
| MCH | 0.1 | 0.0 | 5 | 127 | 80 | 43706 | 0.1313 |
| MCH | 1.0 | 0.0 | 5 | 839 | 792 | 42282 | 1.0697 |
| MHA | 0.01 | 0.0 | 5 | 103 | 8 | 43612 | 0.0706 |
| MHA | 0.1 | 0.0 | 5 | 174 | 79 | 43470 | 0.1614 |
| MHA | 1.0 | 0.0 | 5 | 883 | 788 | 42052 | 1.1019 |
| MHA | 0.01 | 0.2 | 10 | 123 | 28 | 43572 | 0.0961 |
| MHA | 0.01 | 0.3 | 10 | 135 | 40 | 43548 | 0.1114 |
| MHA | 0.01 | 0.4 | 10 | 148 | 53 | 43522 | 0.1281 |
| MHA | 0.01 | 0.5 | 10 | 161 | 66 | 43496 | 0.1447 |

#### Lie-down and recovery times

Event times follow the definitions in Methods, "Orientation, switching and recovery". The median $m_{\mathrm{obs}}$ uses observed events, whereas $m_{\mathrm{clip}}$ includes trajectories without an event at their final observation times. The Kaplan–Meier median $m_{\mathrm{KM}}$ is the first time the estimated survival probability reaches or falls below 0.5. $T$ gives the observation duration or its range across replicates; a dash denotes an unavailable median. For MCH at 0.01 M and +0.1 V, two 2 ns trajectories were too short to satisfy the 5 ns persistence criterion, leaving eight for the event analysis.

Supplementary Table S3: Voltage-driven lie-down times in ns. Medians in Fig. 3L and its associated text use $m_{\mathrm{KM}}$; those in Fig. 3H use $m_{\mathrm{obs}}$. Unreached quantiles in L are displayed at the longest observation time.

| Condition | Events/$n$ | $T$ | $m_{\mathrm{obs}}$ | $m_{\mathrm{clip}}$ | $m_{\mathrm{KM}}$ |
|---|---|---|---|---|---|
| MCH 0.01 M, +0.1 V | 7/8 | 400.0 | 154.10 | 154.95 | 154.10 |
| MCH 0.01 M, +0.2 V | 10/10 | 100.0 | 24.45 | 24.45 | 22.10 |
| MCH 0.01 M, +0.3 V | 10/10 | 100.0 | 10.35 | 10.35 | 9.10 |
| MCH 0.01 M, +0.4 V | 10/10 | 50.0 | 7.65 | 7.65 | 7.30 |
| MCH 0.01 M, +0.5 V | 10/10 | 100.0 | 4.20 | 4.20 | 3.80 |
| MCH 0.15 M, +0.1 V | 5/10 | 400.0 | 222.50 | 382.85 | 365.70 |
| MCH 0.15 M, +0.2 V | 10/10 | 92.8–400.0 | 116.75 | 116.75 | 95.00 |
| MCH 0.15 M, +0.3 V | 10/10 | 200.0 | 32.15 | 32.15 | 30.40 |
| MCH 0.15 M, +0.4 V | 10/10 | 100.0 | 28.10 | 28.10 | 27.90 |
| MCH 0.15 M, +0.5 V | 10/10 | 50.0–400.0 | 29.35 | 29.35 | 28.80 |
| MCH 1.0 M, +0.1 V | 3/10 | 400.0 | 253.40 | 400.00 | – |
| MCH 1.0 M, +0.2 V | 4/10 | 400.0 | 192.70 | 400.00 | – |
| MCH 1.0 M, +0.3 V | 5/10 | 400.0 | 178.80 | 337.00 | 274.00 |
| MCH 1.0 M, +0.4 V | 8/10 | 400.0 | 98.05 | 149.10 | 107.40 |
| MCH 1.0 M, +0.5 V | 7/10 | 100.0–400.0 | 239.00 | 269.90 | 266.20 |
| MHA 0.01 M, +0.2 V | 0/10 | 300.0 | – | 300.00 | – |
| MHA 0.01 M, +0.3 V | 0/10 | 300.0 | – | 300.00 | – |
| MHA 0.01 M, +0.4 V | 6/10 | 300.0–600.0 | 179.45 | 300.00 | 405.60 |
| MHA 0.01 M, +0.5 V | 9/10 | 300.0–600.0 | 109.10 | 130.95 | 109.10 |

Supplementary Table S4: Recovery times after voltage removal or reversal, defined by sustained return to $\theta \leq 75°$. Times are in ns; the main text reports $m_{\mathrm{KM}}$.

| Condition | Events/$n$ | $T$ | $m_{\mathrm{obs}}$ | $m_{\mathrm{clip}}$ | $m_{\mathrm{KM}}$ |
|---|---|---|---|---|---|
| MCH 0.01M, 0 V (75°) | 10/10 | 200.0 | 5.50 | 5.50 | 4.20 |
| MCH 0.15M, 0 V (75°) | 10/10 | 200.0 | 26.50 | 26.50 | 22.20 |
| MCH 1.0M, 0 V (75°) | 5/5 | 200.0 | 66.60 | 66.60 | 66.60 |
| AHT −0.1V, 0.1 M (75°) | 0/3 | 200.0 | – | 200.00 | – |
| AHT −0.2V, 0.1 M (75°) | 0/3 | 200.0 | – | 200.00 | – |
| AHT −0.3V, 0.1 M (75°) | 1/3 | 200.0 | 115.50 | 200.00 | – |
| Bare Au −0.1V, 0.1 M (75°) | 0/3 | 200.0 | – | 200.00 | – |
| Bare Au −0.2V, 0.1 M (75°) | 0/3 | 200.0 | – | 200.00 | – |
| Bare Au −0.3V, 0.1 M (75°) | 0/3 | 200.0 | – | 200.00 | – |
| AHT −0.4V, 0.1 M (75°) | 3/3 | 200.0 | 19.70 | 19.70 | 19.70 |
| AHT −0.5V, 0.1 M (75°) | 3/3 | 200.0 | 4.00 | 4.00 | 4.00 |
| Bare Au −0.4V, 0.1 M (75°) | 0/3 | 200.0 | – | 200.00 | – |
| Bare Au −0.5V, 0.1 M (75°) | 0/3 | 200.0 | – | 200.00 | – |

#### DNA elastic moduli and salt-dependent twist

Present MD values are for 24-bp DNA tethered to MCH-passivated gold at zero bias and 310 K. Moduli at 0.1 M added NaCl use identical frames and fragment lengths sampled within bp 5–21 (interior) or bp 1–24 (end-inclusive).

Supplementary Table S5: Comparison of stretch ($S$), bending ($B$) and torsional ($C$) moduli (Fig. 1F), and salt-dependent helical twist (Fig. 1C), with published experimental and simulation values for DNA in solution. $S$ is in nN; $B$ and $C$ are in nN nm$^2$.

| Observable | Published benchmark | Present simulations |
|---|---|---|
| Salt-dependent helical twist | Experiment: increase of approximately 0.3°/bp from 0.05 to 1.0 M NaCl (7.9-kbp DNA, 295 K, 0.5 pN).[30]<br>Published MD: twist increases from 0.05 to 1.0 M NaCl (25-bp DNA in solution, 295 K).[31] | Mean helical twist increases by 5.2% from 0.01 to 1.0 M added NaCl. |
| Stretch modulus $S$ | Experiment: 0.884 ± 0.116 at 100 mM $Na^+$ (295.65 K);[33] 1.450 ± 0.050 at 50 mM NaCl (293.15 K).[34] | Interior: 1.294 ± 0.202<br>End-inclusive: 0.475 ± 0.060 |
| Bending modulus $B$ | Experiment: 0.196 ± 0.008 at 100 mM $Na^+$ (295.65 K);[33] 0.2223 ± 0.0014 at 100 mM added NaCl (298.15 K).[35] | Interior: 0.222 ± 0.005<br>End-inclusive: 0.0884 ± 0.0165 |
| Torsional modulus $C$ | Experiment: 0.410 ± 0.030 from torque–twist slopes and 0.440 ± 0.040 from twist fluctuations at 100 mM NaCl.[36]<br>Published MD: approximately 0.470 after diameter relaxation at 1.0 M NaCl (25-bp DNA in solution, 295 K).[31] | Interior: 0.469 ± 0.023<br>End-inclusive: 0.0317 ± 0.0104 |

MD moduli are means ± SEM across five independent trajectories. Experimental uncertainty definitions are given in Supplementary Table S6; bending moduli were converted using $B = k_B T l_p$. Twist comparisons are qualitative because the systems, salt ranges and twist definitions differ.

### DNA elasticity in solution

Experimental values used in Fig. 1F are summarized in Supplementary Table S6, with salt conditions and uncertainties as reported in the original studies. Forth and Kriegel report effective torsional stiffnesses under tension. The Kriegel band represents the reported high-force result over 0.02–0.5 M NaCl.

Supplementary Table S6: Experimental DNA elastic properties used in Fig. 1F. Stretch modulus $S$ is in pN, bending persistence length $l_p$ in nm and torsional modulus $C$ in pN nm$^2$, unless indicated.

| Source | Condition | Reported value | Measurement and uncertainty |
|---|---|---|---|
| Wenner 2002 [33] | 10 mM $Na^+$ | $S$ = 741 ± 147; $l_p$ = 67 ± 4 | Mean ± SE, $n \geq 3$; three-parameter fit. |
| Wenner 2002 | 100 mM $Na^+$ | $S$ = 884 ± 116; $l_p$ = 48 ± 2 | Mean ± SE; same fit as above. |
| Wenner 2002 | 1000 mM $Na^+$ | $S$ = 1256 ± 217; $l_p$ = 46 ± 2 | Mean ± SE; same fit as above. |
| Wang 1997 [46] | 10 mM $Na^+$, phosphate buffer | $S$ = 1008 ± 38; $l_p$ = 47.4 ± 1.0 | Mean ± SE, $n$ = 10 for $S$ and $n$ = 14 for $l_p$. |
| Bustamante 1994 [47] | 97-kbp $\lambda$-DNA dimer | $l_p$ = 53.4 ± 2.3 | 90% confidence interval from a WLC fit. |
| Guilbaud 2019 [35] | 100 mM added NaCl; $I$ = 100.5 mM | $l_p$ = 54.01 ± 0.33 | Bootstrap SD of $l_p$. |
| Baumann 1997 [32] | $I$ = 1.86 mM, monovalent buffer | $S$ = 649 ± 82; $l_p$ = 94.9 ± 5.9 | Mean ± SE, $n \geq 3$; extensible-WLC fit. |
| Baumann 1997 | $I$ = 586 mM | $S$ = 1435 ± 160; $l_p$ = 55.9 ± 3.2 | Mean ± SE; all nine salt conditions are included in Fig. 1F. |
| Gross 2011 [34] | 50 mM NaCl, 10 mM Tris-HCl; 20 °C | $S$ = 1450 ± 50; $l_p$ = 38 ± 2 | Mean ± SEM, seven molecules; twistable-WLC fit. |
| Bryant 2003 [36] | 100 mM NaCl | $C$ = 410 ± 30 | Torque–twist slope; torque calibration dominates the uncertainty. |
| Bryant 2003 | Same buffer | $C$ = 440 ± 40 | Twist fluctuations at 15 pN, five molecules. |
| Mosconi 2009 [37] | 100 mM NaCl | $C/k_BT$ = 94 ± 7 nm | Intrinsic limit from the Moroz–Nelson correction; uncertainty type unspecified. |
| Forth 2008 [48] | 150 mM NaCl in phosphate-buffered saline; 23.5 °C | $C_{\mathrm{eff}}/k_BT$ = 90±3 and 88±4 nm; $l_p$ = 43±3 | Torque slopes for 2.2- and 4.2-kbp DNA; uncertainty type unspecified. |
| Kriegel 2017 [49] | Multiple buffers; 6.5 pN | $C_{\mathrm{eff}}/k_BT$ = 103 ± 4 nm | High-force result for 20–500 mM NaCl and 100 mM NaCl with 10 mM $MgCl_2$. |

Bending moduli were calculated as $B = k_{\mathrm{B}}Tl_p$. We used 295.65 K for Wenner, 293.15 K for Gross, 296.65 K for Forth and 298.15 K for Wang, Bustamante, Guilbaud and Baumann. The temperatures for Wang and Bustamante were assumed. Torsional persistence lengths from Mosconi and Kriegel were converted at 298.15 K. The 10 mM $Na^+$ condition for Bustamante follows Marko and Siggia (*Macromolecules* 1995, 28, 8759–8770).
